# OmicSelector: automatic feature selection and deep learning modeling for omic experiments

**DOI:** 10.1101/2022.06.01.494299

**Authors:** Konrad Stawiski, Marcin Kaszkowiak, Damian Mikulski, Piotr Hogendorf, Adam Durczyński, Janusz Strzelczyk, Dipanjan Chowdhury, Wojciech Fendler

## Abstract

A crucial phase of modern biomarker discovery studies is selecting the most promising features from high-throughput screening assays. Here, we present the OmicSelector - Docker-based web application and R package that facilitates the analysis of such experiments. OmicSelector provides a consistent and overfitting-resilient pipeline that integrates 94 feature selection approaches based on 25 distinct variable selection methods. It identifies and then ranks the best feature sets using 11 modeling techniques with hyperparameter optimization in hold-out or cross-validation. OmicSelector provides classification performance metrics for proposed feature sets, allowing researchers to choose the overfitting-resistant biomarker set with the highest diagnostic potential. Finally, it performs GPU-accelerated development, validation, and implementation of deep learning feedforward neural networks (up to 3 hidden layers, with or without autoencoders) on selected signatures. The application performs an extensive grid search of hyperparameters, including balancing and preprocessing of next-generation sequencing (e.g. RNA-seq, miRNA-seq) oraz qPCR data. The pipeline is applicable for determining candidate circulating or tissue miRNAs, gene expression data and methylomic, metabolomic or proteomic analyses. As a case study, we use OmicSelector to develop a diagnostic test for pancreatic and biliary tract cancer based on serum small RNA next-generation sequencing (miRNA-seq) data. The tool is open-source and available at https://biostat.umed.pl/OmicSelector/

## 1. Introduction

Broad-scale treatment personalization is one of the most significant modern medicine challenges, requiring accurate and cost-effective diagnostic tests. Such methods rely heavily on biomarkers, which are usually discovered using *omic* techniques. Although high-throughput experiments accumulate extensive amount of biomarker candidates, translating the results to the clinical bedside remains troublesome.

The typical biomarker study consists of discovery and validation phases. [1] In the former, high-throughput screening is usually performed to measure the values of multiple features which are further assessed to determine their diagnostic potential. Afterwards, only the best ones are measured in the validation phase, typically in a new set of samples, with a cheaper and/or more accessible method.

Our team has been working on microRNA (miRNA) biomarkers for radiation [2] and cancer [3], but identified issues with the reproducibility of selected biomarker performance [4–6] or reference identification [7] that hinder direct translation of experimental models to clinical medicine. Similar challenges, caused by bias and overfitting, hindered other groups’ attempts to develop validated, efficient omic-driven biomarkers. [8]

Cohorts used in the discovery phase of biomarker studies are usually small due to the high cost of high-throughput assays, which makes the experiments vulnerable to overfitting and results in false-positive biomarker candidates that fail in external validation. [9] Recent reviews and meta-analyses of serum miRNA biomarkers for pancreatic cancer [10,11] highlight how various miRNA sets are chosen in different studies, all of which report unrealistically optimistic results. Noteworthy, some miRNAs were frequently selected as references for internal standardization of specific assays, but, at the same time, other authors report their significant dysregulation in pancreatic cancer cases. For example, Johansen et al. [12] show hsa-miR-16 to be significantly upregulated in pancreatic cancer, while several other papers use this miRNA as stable reference. [13,14] Lack of external validation further hinders efficient diagnostic test development. Thus, correct and overfitting-resistant feature selection is critical in biomarker studies.

Recent years have also provided significant progress in the development of classification models. [15] Although deep learning [16] can provide a meaningful boost in classifiers’ performance, its power is rarely applied to common laboratory or clinical data due to technical limitations and threshold knowledge.

In this paper, we try to tackle this problem by designing and implementing software for systematic, overfitting-resistant, and informative feature selection with subsequent development of the deep learning-based diagnostic model.

## 2. Implementation

The analytical steps of our package entail (Figure 1): splitting of the dataset into training, testing and validation sets, differential expression analysis, performing up to 94 different feature selection procedures on the training set, testing of feature sets (formulas), and deep learning model development. Formulas are validated in terms of their potential of creating efficient classification models by training 11 models of various approaches with hyperparameter optimization based on hold-out- or cross-validation. After the feature set is chosen, OmicSelector performs extensive deep feedforward neural network modeling with hyperparameter optimization based on grid search. The software searches for the best neural network up to 3 hidden layers and with or without additional feature extraction by (sparse) autoencoders. (Figure 2) Our toolset enables the users to make an informed decision about the most appropriate feature selection method, informs them about their predictive abilities using different modeling approaches, and develops a final deep neural network that can be efficiently utilized for further predictions on new data.

**Figure 1.**
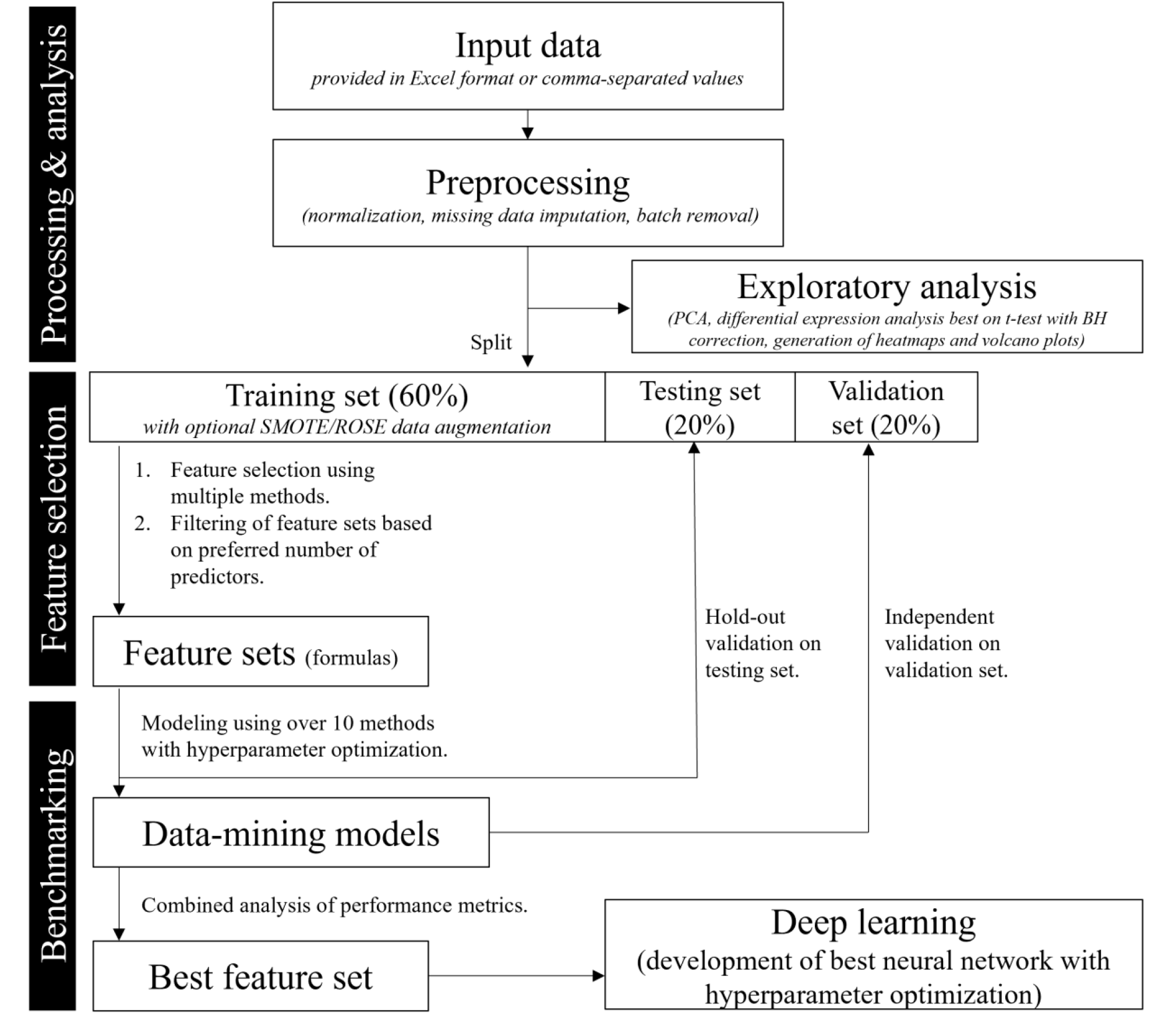
The pipeline of OmicSelector analysis. Abbreviations: PCA – principal component analysis, BH - Benjamini-Hochberg procedure, SMOTE/ROSE – data balancing methods explained in the main text.

**Figure 2.**
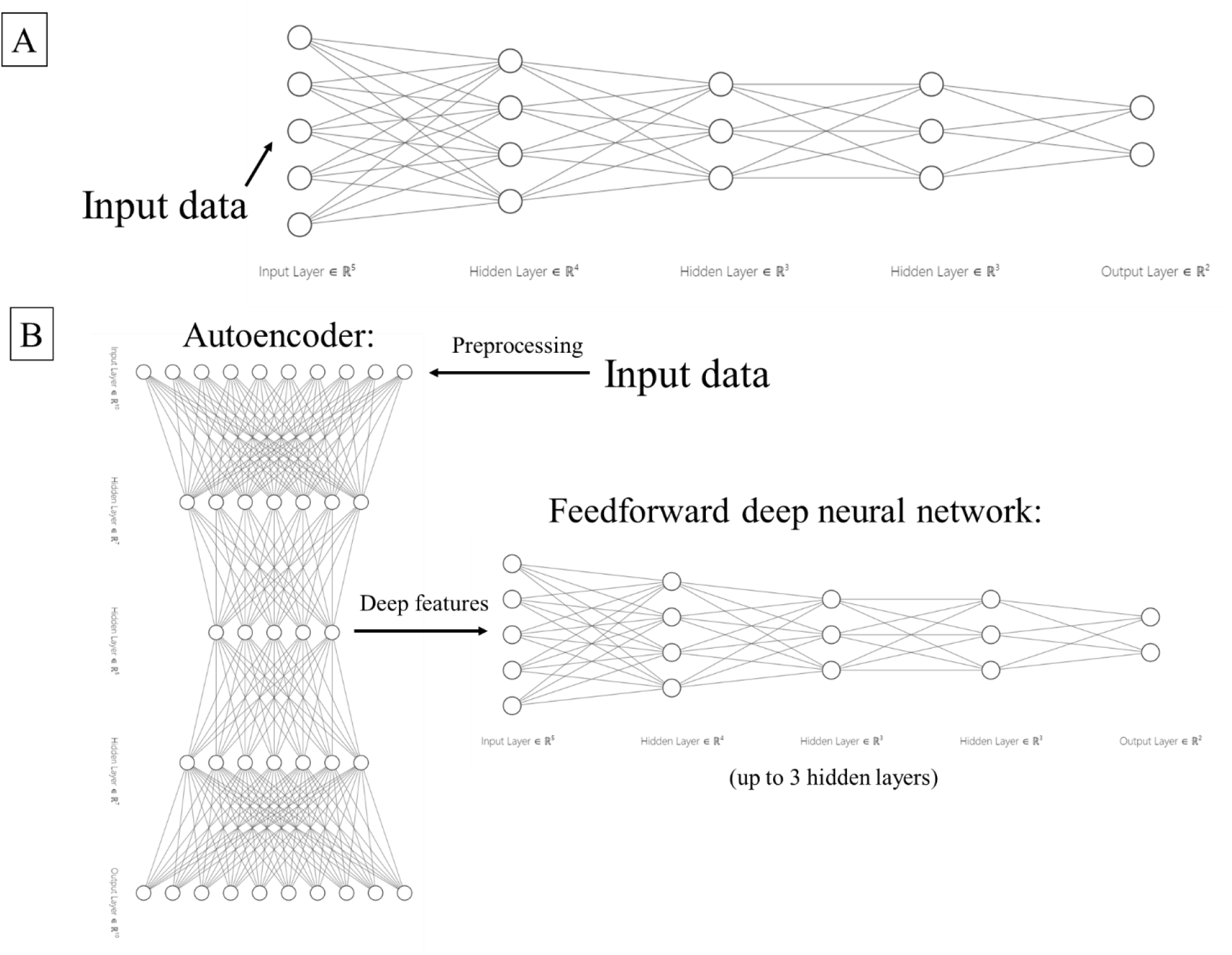
Example structure of deep neural networks trained by OmicSelector. Panel A shows a feedforward neural network up to 3 hidden layers. In this scenario, when no autoencoder is used, input data (after preprocessing) are provided to the network’s input layer. Panel B shows the exemplary structure of networks when an autoencoder is applied. First, the input data (after preprocessing) are processed in an autoencoder. In the next step, deep features are extracted from the bottleneck layer and serve as an input in a feedforward deep neural network. Sparsity can be implemented in autoencoders using L1 regularization.

OmicSelector is an extensive R package with a PHP-based web graphical user interface(GUI) distributed with its environment confined as a Docker image.

Crude or log10-transformed expression values can be used as an input in the analysis. The dataset is then randomly split into training, testing, or validation sets in default percentages of: 60%:20%:20% respectively or according to the user’s split predefined. The exploratory analysis is composed of principal component and differential expression analysis. Expression heatmaps with clustering and volcano plots are generated to visualize the data and allow the user to identify outliers, and/or batch effects. Our package can also efficiently perform normalization of sequencing read counts, correcting the batch effect using Combat [17] or imputing missing data with predictive mean matching or using the mean or median expression. [18] Due to potential problems with efficient classifier development associated with imbalanced datasets, the software allows the user to attempt to balance the number of cases and controls by using methods such as SMOTE [19] or ROSE [20].

As a core functionality, our software integrates 94 feature selection approaches described in detail in Table 1. Briefly, we use 25 feature selection methods or software packages and apply them to the whole dataset, a prefiltered dataset with features that are significant in differential analysis, and on balanced datasets (both whole and prefiltered).

**Table 1.**
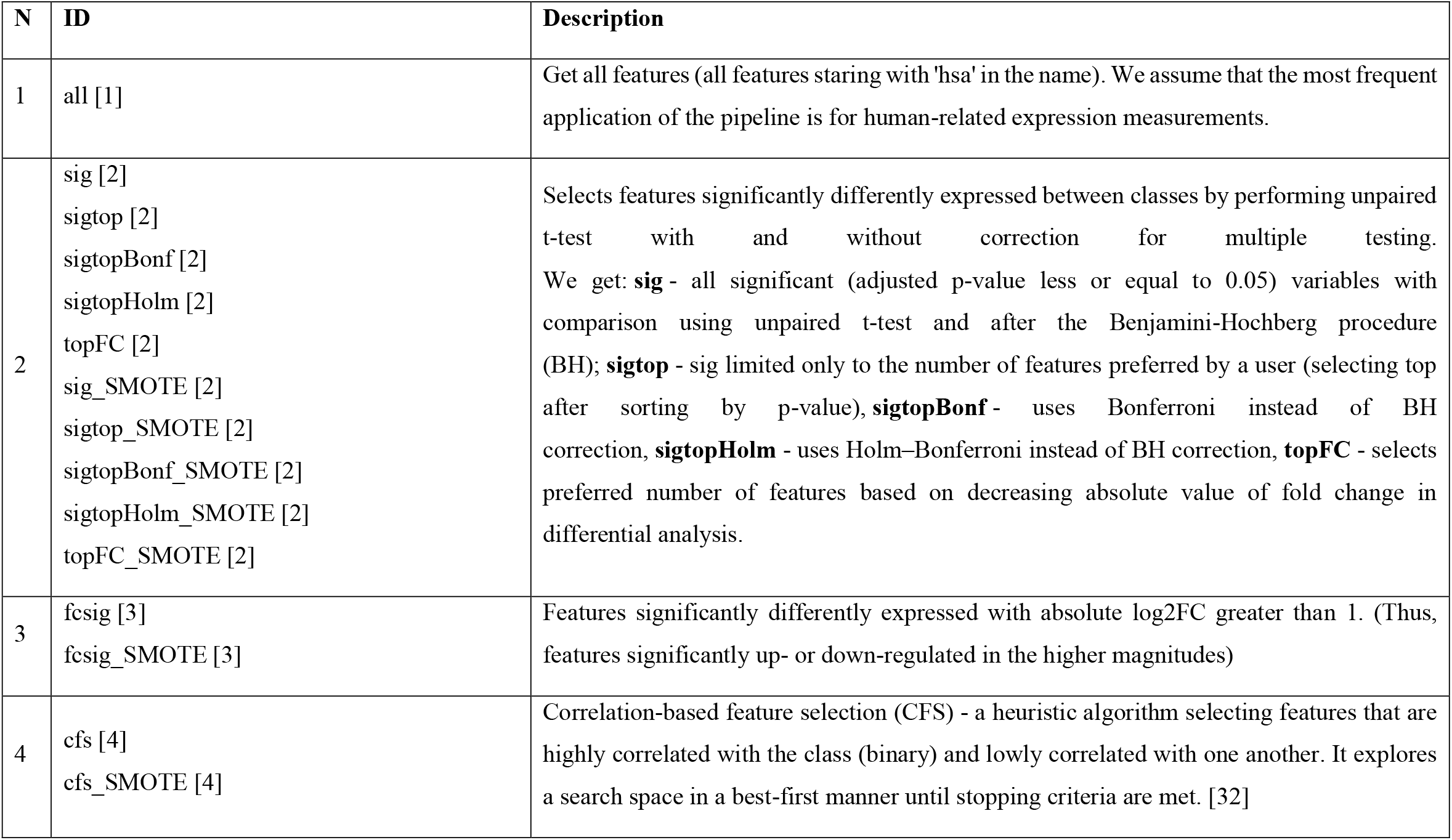
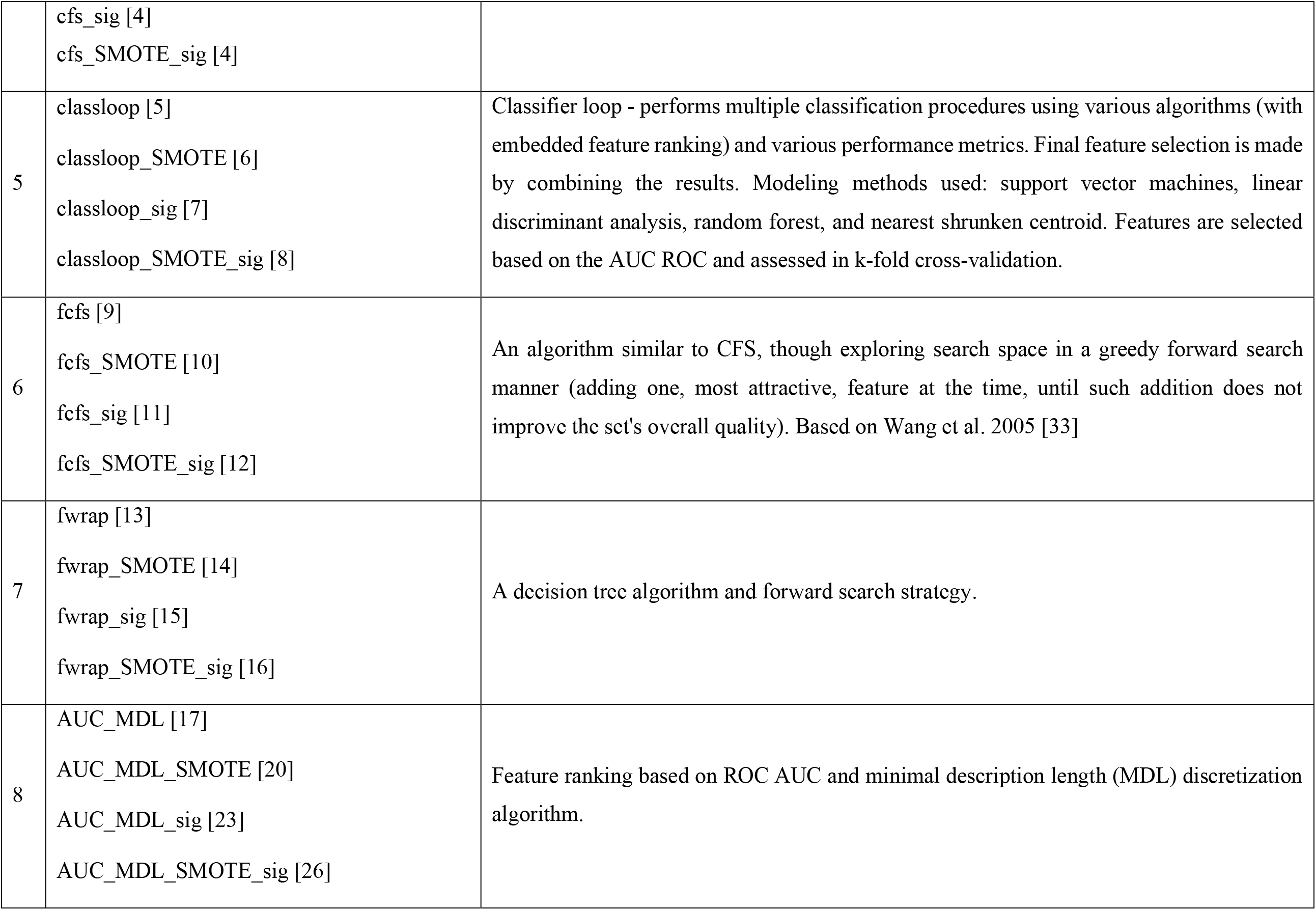
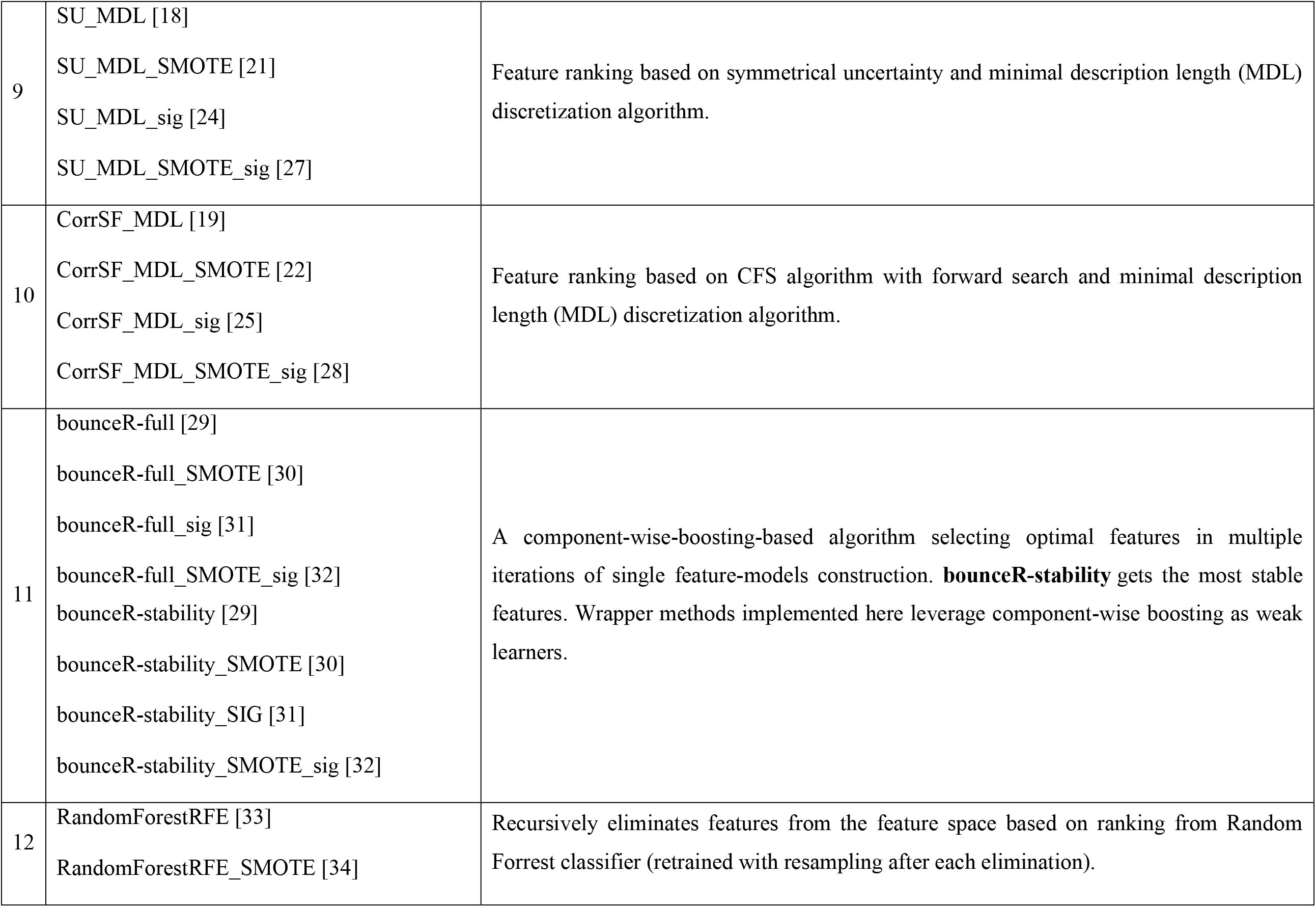
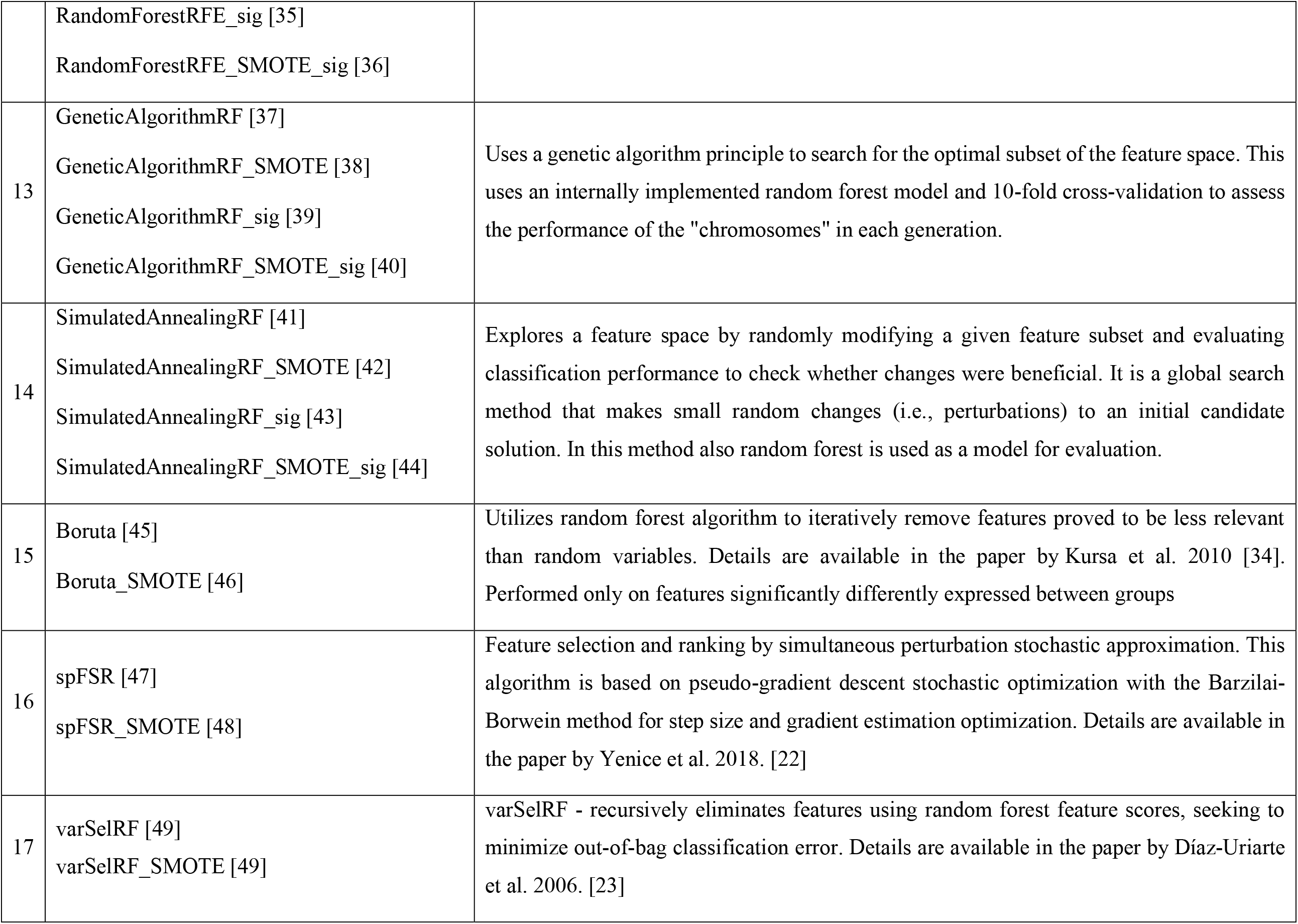
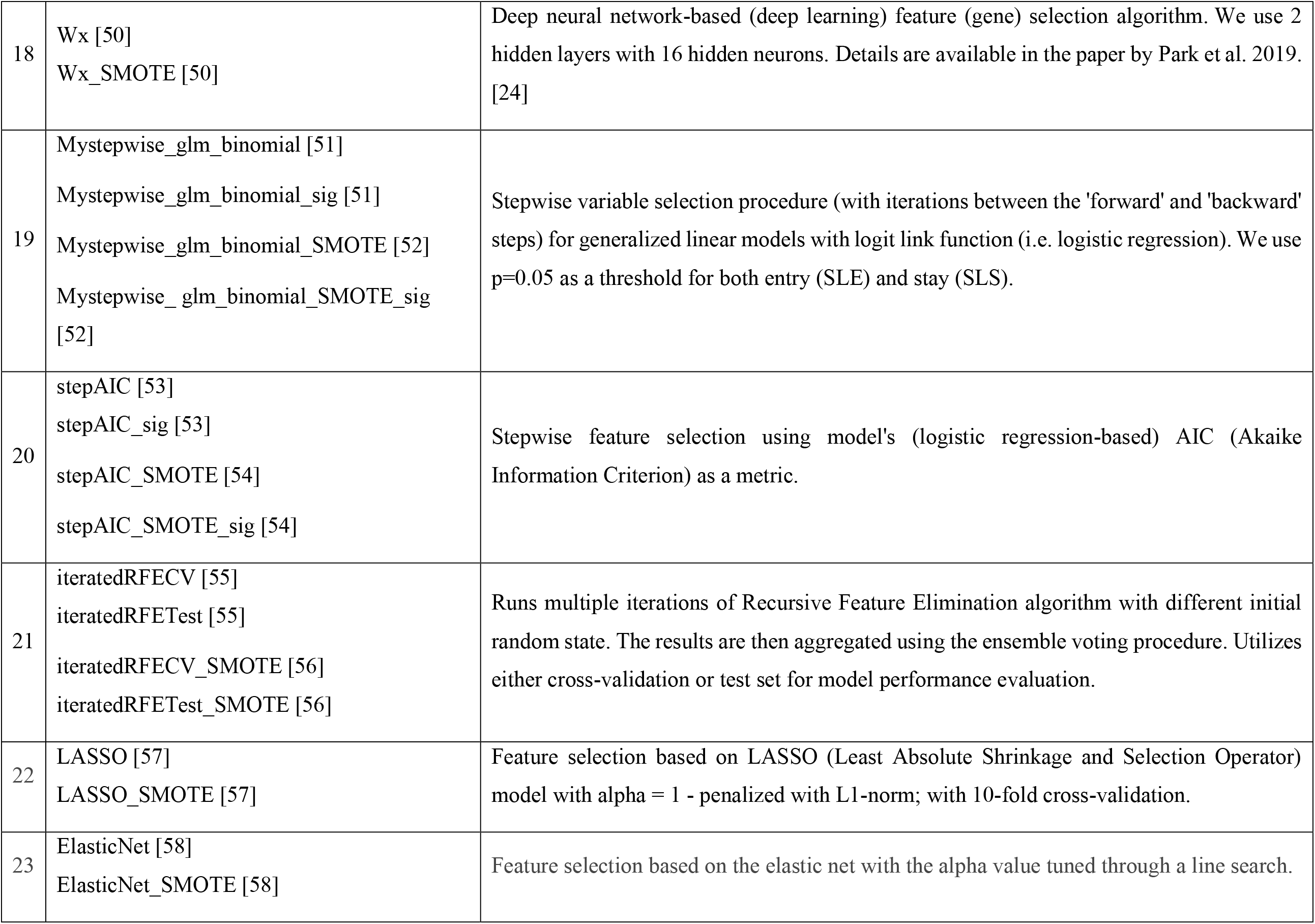
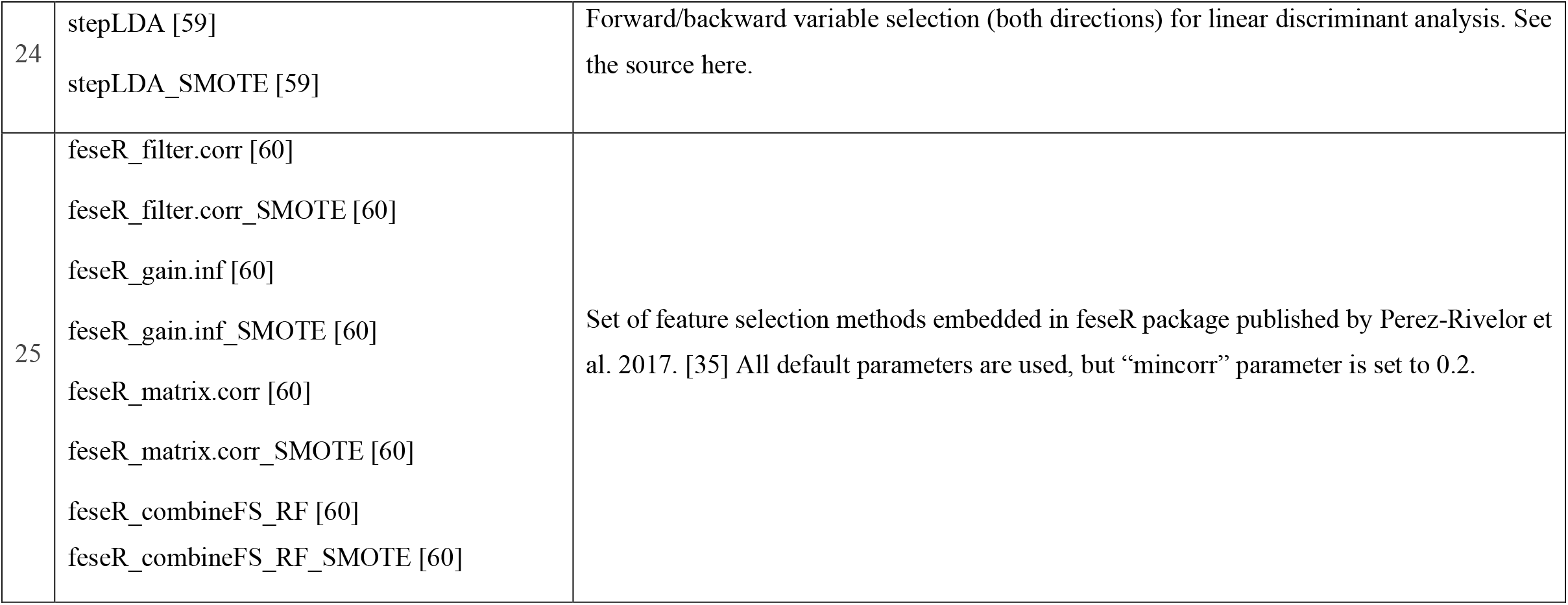
Feature selection methods available in the package. Selected methods are also performed on datasets augmented with Synthetic Minority Oversampling Technique (SMOTE) [19] – their names are then appended with “_SMOTE”. The majority of methods can be performed on a whole feature space or only on the subset of features significantly differently expressed between groups (indicated with a “_sig” suffix). The values in brackets indicate the value of the “m” parameter used in *OmicSelector_OmicSelector()* method.

| N | ID | Description |
| --- | --- | --- |
| 1 | all [1] | Get all features (all features starting with 'hsa' in the name). We assume that the most frequent application of the pipeline is for human-related expression measurements. |
| 2 | sig [2]<br>sigtop [2]<br>sigtopBonf [2]<br>sigtopHolm [2]<br>topFC [2]<br>sig_SMOTE [2]<br>sigtop_SMOTE [2]<br>sigtopBonf_SMOTE [2]<br>sigtopHolm_SMOTE [2]<br>topFC_SMOTE [2] | Selects features significantly differently expressed between classes by performing unpaired t-test with and without correction for multiple testing. We get: **sig** - all significant (adjusted p-value less or equal to 0.05) variables with comparison using unpaired t-test and after the Benjamini-Hochberg procedure (BH); **sigtop** - sig limited only to the number of features preferred by a user (selecting top after sorting by p-value), **sigtopBonf** - uses Bonferroni instead of BH correction, **sigtopHolm** - uses Holm–Bonferroni instead of BH correction, **topFC** - selects preferred number of features based on decreasing absolute value of fold change in differential analysis. |
| 3 | fcsig [3]<br>fcsig_SMOTE [3] | Features significantly differently expressed with absolute log2FC greater than 1. (Thus, features significantly up- or down-regulated in the higher magnitudes) |
| 4 | cfs [4]<br>cfs_SMOTE [4] | Correlation-based feature selection (CFS) - a heuristic algorithm selecting features that are highly correlated with the class (binary) and lowly correlated with one another. It explores a search space in a best-first manner until stopping criteria are met. [32] |
|  | cfs_sig [4]<br>cfs_SMOTE_sig [4] |  |
| 5 | classloop [5]<br>classloop_SMOTE [6]<br>classloop_sig [7]<br>classloop_SMOTE_sig [8] | Classifier loop - performs multiple classification procedures using various algorithms (with embedded feature ranking) and various performance metrics. Final feature selection is made by combining the results. Modeling methods used: support vector machines, linear discriminant analysis, random forest, and nearest shrunken centroid. Features are selected based on the AUC ROC and assessed in k-fold cross-validation. |
| 6 | fcfs [9]<br>fcfs_SMOTE [10]<br>fcfs_sig [11]<br>fcfs_SMOTE_sig [12] | An algorithm similar to CFS, though exploring search space in a greedy forward search manner (adding one, most attractive, feature at the time, until such addition does not improve the set's overall quality). Based on Wang et al. 2005 [33] |
| 7 | fwrap [13]<br>fwrap_SMOTE [14]<br>fwrap_sig [15]<br>fwrap_SMOTE_sig [16] | A decision tree algorithm and forward search strategy. |
| 8 | AUC_MDL [17]<br>AUC_MDL_SMOTE [20]<br>AUC_MDL_sig [23]<br>AUC_MDL_SMOTE_sig [26] | Feature ranking based on ROC AUC and minimal description length (MDL) discretization algorithm. |
| 9 | SU_MDL [18]<br>SU_MDL_SMOTE [21]<br>SU_MDL_sig [24]<br>SU_MDL_SMOTE_sig [27] | Feature ranking based on symmetrical uncertainty and minimal description length (MDL) discretization algorithm. |
| 10 | CorrSF_MDL [19]<br>CorrSF_MDL_SMOTE [22]<br>CorrSF_MDL_sig [25]<br>CorrSF_MDL_SMOTE_sig [28] | Feature ranking based on CFS algorithm with forward search and minimal description length (MDL) discretization algorithm. |
| 11 | bounceR-full [29]<br>bounceR-full_SMOTE [30]<br>bounceR-full_sig [31]<br>bounceR-full_SMOTE_sig [32]<br>bounceR-stability [29]<br>bounceR-stability_SMOTE [30]<br>bounceR-stability_SIG [31]<br>bounceR-stability_SMOTE_sig [32] | A component-wise-boosting-based algorithm selecting optimal features in multiple iterations of single feature-models construction. **bounceR-stability** gets the most stable features. Wrapper methods implemented here leverage component-wise boosting as weak learners. |
| 12 | RandomForestRFE [33]<br>RandomForestRFE_SMOTE [34] | Recursively eliminates features from the feature space based on ranking from Random Forrest classifier (retrained with resampling after each elimination). |
|  | RandomForestRFE_sig [35]<br>RandomForestRFE_SMOTE_sig [36] |  |
| 13 | GeneticAlgorithmRF [37]<br>GeneticAlgorithmRF_SMOTE [38]<br>GeneticAlgorithmRF_sig [39]<br>GeneticAlgorithmRF_SMOTE_sig [40] | Uses a genetic algorithm principle to search for the optimal subset of the feature space. This uses an internally implemented random forest model and 10-fold cross-validation to assess the performance of the "chromosomes" in each generation. |
| 14 | SimulatedAnnealingRF [41]<br>SimulatedAnnealingRF_SMOTE [42]<br>SimulatedAnnealingRF_sig [43]<br>SimulatedAnnealingRF_SMOTE_sig [44] | Explores a feature space by randomly modifying a given feature subset and evaluating classification performance to check whether changes were beneficial. It is a global search method that makes small random changes (i.e., perturbations) to an initial candidate solution. In this method also random forest is used as a model for evaluation. |
| 15 | Boruta [45]<br>Boruta_SMOTE [46] | Utilizes random forest algorithm to iteratively remove features proved to be less relevant than random variables. Details are available in the paper by Kursa et al. 2010 [34]. Performed only on features significantly differently expressed between groups |
| 16 | spFSR [47]<br>spFSR_SMOTE [48] | Feature selection and ranking by simultaneous perturbation stochastic approximation. This algorithm is based on pseudo-gradient descent stochastic optimization with the Barzilai-Borwein method for step size and gradient estimation optimization. Details are available in the paper by Yenice et al. 2018. [22] |
| 17 | varSelRF [49]<br>varSelRF_SMOTE [49] | varSelRF - recursively eliminates features using random forest feature scores, seeking to minimize out-of-bag classification error. Details are available in the paper by Díaz-Uriarte et al. 2006. [23] |
| 18 | W<sub>x</sub> [50]<br>W<sub>x</sub>_SMOTE [50] | Deep neural network-based (deep learning) feature (gene) selection algorithm. We use 2 hidden layers with 16 hidden neurons. Details are available in the paper by Park et al. 2019. [24] |
| 19 | Mystepwise_glm_binomial [51]<br>Mystepwise_glm_binomial_sig [51]<br>Mystepwise_glm_binomial_SMOTE [52]<br>Mystepwise_glm_binomial_SMOTE_sig [52] | Stepwise variable selection procedure (with iterations between the 'forward' and 'backward' steps) for generalized linear models with logit link function (i.e. logistic regression). We use p=0.05 as a threshold for both entry (SLE) and stay (SLS). |
| 20 | stepAIC [53]<br>stepAIC_sig [53]<br>stepAIC_SMOTE [54]<br>stepAIC_SMOTE_sig [54] | Stepwise feature selection using model's (logistic regression-based) AIC (Akaike Information Criterion) as a metric. |
| 21 | iteratedRFECV [55]<br>iteratedRFETest [55]<br>iteratedRFECV_SMOTE [56]<br>iteratedRFETest_SMOTE [56] | Runs multiple iterations of Recursive Feature Elimination algorithm with different initial random state. The results are then aggregated using the ensemble voting procedure. Utilizes either cross-validation or test set for model performance evaluation. |
| 22 | LASSO [57]<br>LASSO_SMOTE [57] | Feature selection based on LASSO (Least Absolute Shrinkage and Selection Operator) model with alpha = 1 - penalized with L1-norm; with 10-fold cross-validation. |
| 23 | ElasticNet [58]<br>ElasticNet_SMOTE [58] | Feature selection based on the elastic net with the alpha value tuned through a line search. |
| 24 | stepLDA [59]<br>stepLDA_SMOTE [59] | Forward/backward variable selection (both directions) for linear discriminant analysis. See the source here. |
| 25 | feseR_filter.corr [60]<br>feseR_filter.corr_SMOTE [60]<br>feseR_gain.inf [60]<br>feseR_gain.inf_SMOTE [60]<br>feseR_matrix.corr [60]<br>feseR_matrix.corr_SMOTE [60]<br>feseR_combineFS_RF [60]<br>feseR_combineFS_RF_SMOTE [60] | Set of feature selection methods embedded in feseR package published by Perez-Rivelor et al. 2017. [35] All default parameters are used, but “mincorr” parameter is set to 0.2. |

The feature selection methods include classical ones based on significance in differential analysis, correlation-based feature selection methods, stepwise modeling approaches, feature ranking based on different metrics and minimal description length discretization algorithm, and recursive feature elimination. However, more advanced and recent methods like component-wise-boosting-based algorithms, genetic algorithms, simulated annealing, Boruta [21], spFSR [22], varSelRF [23] or deep neural network-based WxNet [24] are also implemented. The user can set the maximum number of features, which are further selected based on the training set data.

The selected feature sets are evaluated (i.e., “benchmarked”) by introducing various classification models. The training set is used for model development, which is further optimized (hyperparameter tuning with random search) by evaluation of its performance on the test set (by default; as shown in Figure 1) or in repeated cross-validation (5 times 10-fold cross-validation). This step’s primary purpose is to find the best set of features, which is resilient to overfitting.

In Table 2 we presented the methods used during benchmarking. Those include a broad spectrum of approaches starting from simple logistic regression and decision trees and ending with deep neural networks. The final models are scored on this independent validation set. By default, the pipeline chooses the best signature based on the highest harmonic mean of training, testing, and validation accuracies in all models, i.e.:

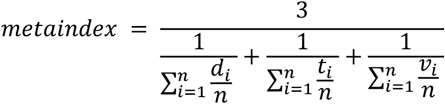

where: *d* – accuracy on the training set, *t* – accuracy on the testing set, *v* – accuracy on the validation set, *n* – number of feature selection procedures. The application of harmonic mean ensures the equal contribution of all three datasets. The final analysis of selected signatures allows us to visualize sets of features across the entire dataset and highlight a classification potential.

**Table 2.** Modeling methods available in the package. ID is the codename used in the software.

| ID | Description |
| --- | --- |
| glm | Logistic regression (generalized linear model with binomial link function). |
| mlp | Multilayer perceptron (MLP) - fully connected feedforward neural network with 1 hidden layer and logistic activation function. |
| mlpML | Multilayer perceptron (MLP) - fully connected feedforward neural network with up to 3 hidden layers and logistic activation function. |
| svmRadial | Support vector machines with radial basis function kernel. |
| svmLinear | Support vector machines with a linear kernel. |
| rf | Random forest model. |
| C5.0 | C5.0 decision trees and rule-based models. |
| rpart | CART decision trees with modulation of complexity parameter. |
| rpart2 | CART decision trees with modulation of max tree depth. |
| ctree | Conditional inference trees. |
| xgbTree | eXtreme gradient boosting. |

OmicSelector performs extensive GPU-accelerated deep feedforward neural network modeling with hyperparameter optimization based on grid search. Proposed software inducts neural networks up to 3 hidden layers and with or without additional feature extraction by (sparse) autoencoders and offers different approaches depending on available computational power.

Standard full scan (without autoencoders) trains 97848 neural networks with different hyperparameters. It can be further extended by including autoencoders, which cause OmicSelector to train up to 293544 neural networks in the extended scan. Hyperparameters tuned in the standard full scan were shown in Figure 3. The set of hyperparameters for the extended scan was provided in Supplementary Table 1.

**Figure 3.**
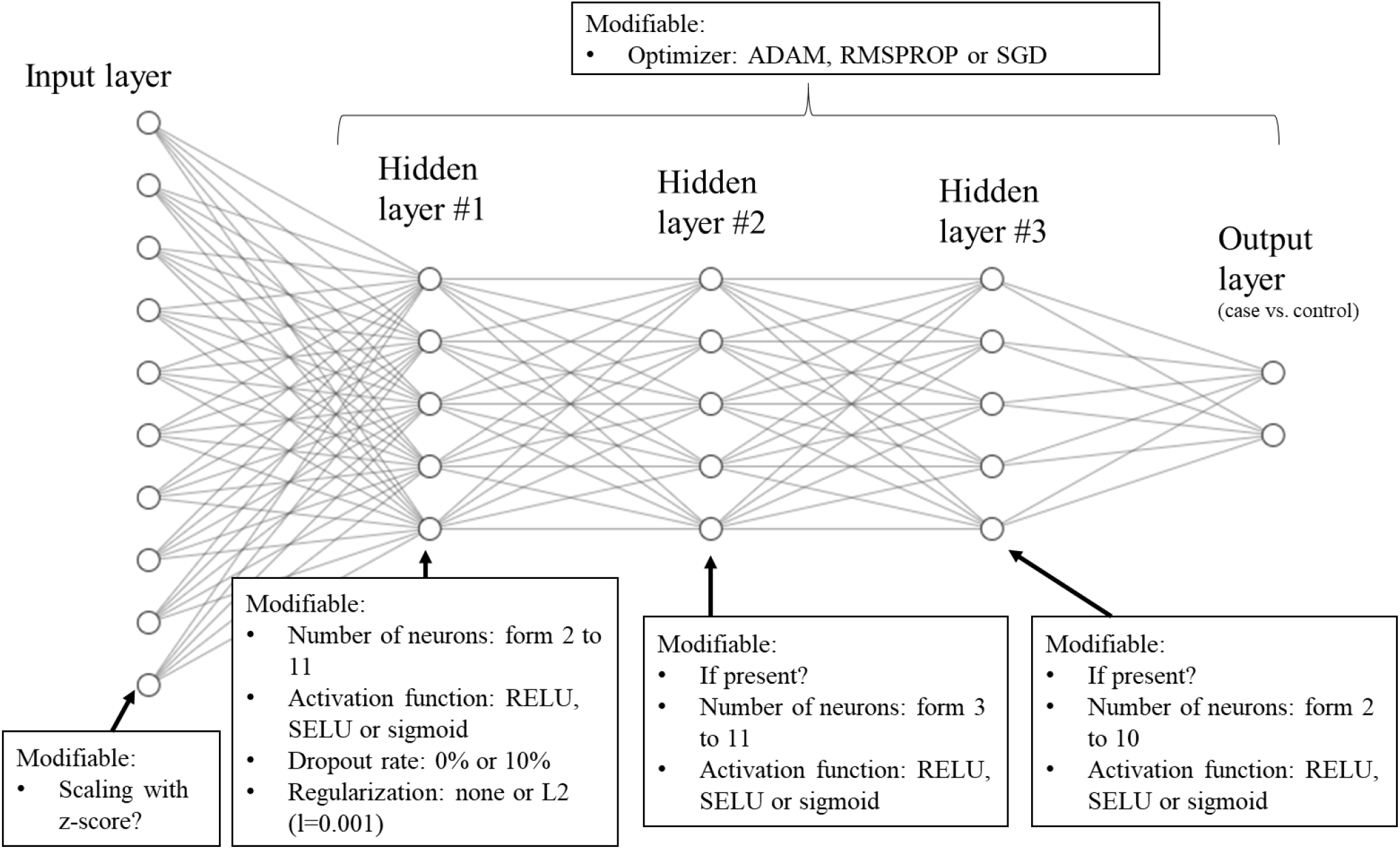
Hyperparameters tuned in artificial neural networks developed in a standard full scan of OmicSelector.

Training parameters are described in much detail in the official documentation on the project website. Briefly, feedforward neural networks (without autoencoders) are trained for 2000 epochs with early stopping. Before training, apart from balancing, the dataset can also be scaled (using a z-score). Scaling parameters are calculated on the training set and reapplied on testing and validation sets. The induction is conducted on the training set with standard concurrent loss analysis on the testing set (but not the validation set). If no improvement in testing loss is seen for 50 epochs, the training stops. This step allowed for controlling the training process before reaching overfitting, thus increasing the testing set’s error. The final model with the lowest testing error is chosen by the analysis of loss fluctuations in training history. Therefore, the selected neural network for further investigation is the latest network with the lowest error on the testing set. Importantly, this means that the newest network in the training process is chosen, and the early stopping cannot occur before the 50^th^ epoch, so the chance of selecting an untrained network with overstated testing performance is minimized.

Based on receiver operating curve analysis (ROC) and the Youden index, the best cutoff value for the predicted probability of the final neural network is estimated on the training set and used in performance assessment on testing and validation sets. The best model is finally chosen based on the highest model metaindex value. This metric is defined as the harmonic average of all accuracy metrics (on training, testing, and validation sets). It is, therefore, identical to the metaindex presented above with parameter *n* equal to 1 (*n*=1).

For some hyperparameter combinations, the autoencoder-based deep feature extraction can be added to the network structure. The structural differences between the networks with and without 5-layer autoencoders were shown in Figure 2. Additional sparsity can be introduced to autoencoders with L1 regularization (L1 regularization penalty of 0.001).

## 3. Results

The OmicSelector was developed, tested, and implemented as a Docker-based application. The source code was placed on GitHub, and continuous integration was set up to deliver the newest builds to Docker Hub directly. The docker-based application provides a PHP-based web interface to the R package and additional tools: R Shiny applications for preprocessing and model implementation; integrated development environment with R Studio, Jupyter-based notebooks, and VS Code. Every analysis can be further extended with R or python code directly in the application.

The R package can be built directly in new environments, or the package can be quickly installed with its dependencies using conda-pack. The publicly available free version based on our application’s prebuilt Docker image was provided using the Ainize service.

Website, documentation, tutorials, and demo version were provided at https://biostat.umed.pl/OmicSelector/

### 3.1 Case study

In 2019, Kim et al. [25] published a small RNA next-generation sequencing (miRNA-seq) serum study detailing the expression profiles of 34 patients with pancreatic (PCa) or biliary tract cancers (BTCa) and 21 controls. Based on measurements of 677 miRNAs detected in blood samples, the authors suggested that hsa-miR-744-5p, hsa-miR-409-3p, and hsa-miR-128-3p could serve as potential biomarkers for PCa and BTCa diagnosis. The authors found that three of the best performing miRNAs could distinguish cancer patients from controls, with an accuracy of 92.7%. The data from miRNA-seq were deposited in GEO with accession number GSE109319.

To check whether OmicSelector could provide a more robust signature, we applied our pipeline to the referenced set. After running all feature selection methods, only features with less than eleven miRNAs were included in further steps. The signature selected by the authors of the original report (hsa-miR-744-5p, hsa-miR-409-3p, and hsa-miR-128-3p) was manually added to established formulas before benchmarking. Default OmicSelectors parameters and methods were used in the evaluation of signatures. After the best set of miRNAs was selected based on the highest benchmarking metaindex values, we conducted a full deep learning scan (97848 deep feedforward neural networks with diverse hyperparameters) on both the signature selected by OmicSelector and the one by authors of the original publication. Supplementary Table 2 presents sets of hyperparameters assessed in this scan. The general performance and superiority of methods/hyperparameters were evaluated using a linear mixed-effects model with varying intercepts for each hyperparameter setting. Model metaindex was used as a dependent variable, and hyperparameters were treated as predictors. Model estimates and their 95%CI were used in statistical inference. Average model metaindex values were compared using an unpaired t-test.

Data downloaded from GEO were correctly formatted and uploaded to OmicSelector. After feature selection and filtering, the final 53 signatures proposed by OmicSelector were subjected to benchmarking. (Supplementary Table 3) The three best-performing signatures were:

1. “cfs” – hsa-miR-125b-5p, hsa-miR-148b-3p, hsa-miR-223-5p, hsa-miR-361-5p, hsa-miR-4433b-5p, and hsa-miR-744-5p (metaindex of 0.8809)
2. “ElasticNet_SMOTE” – hsa-miR-744-5p, hsa-miR-128-3p, hsa-miR-125b-5p, hsa-miR-152-3p, hsa-miR-142-5p, hsa-miR-148b-3p, hsa-miR-1307-3p, hsa-miR-24-3p, hsa-miR-222-3p, and hsa-miR-122-5p (metaindex of 0.8702)
3. “Original” (Authors’ signature) – hsa-miR-744-5p, hsa-miR-409-3p, and hsa-miR-128-3p (metaindex of 0.8686)

The overlap of those signatures is shown in Figure 4A. Metrics for individual models in benchmarking were provided in Supplementary Table 4 and Supplementary Table 5.

**Figure 4.**
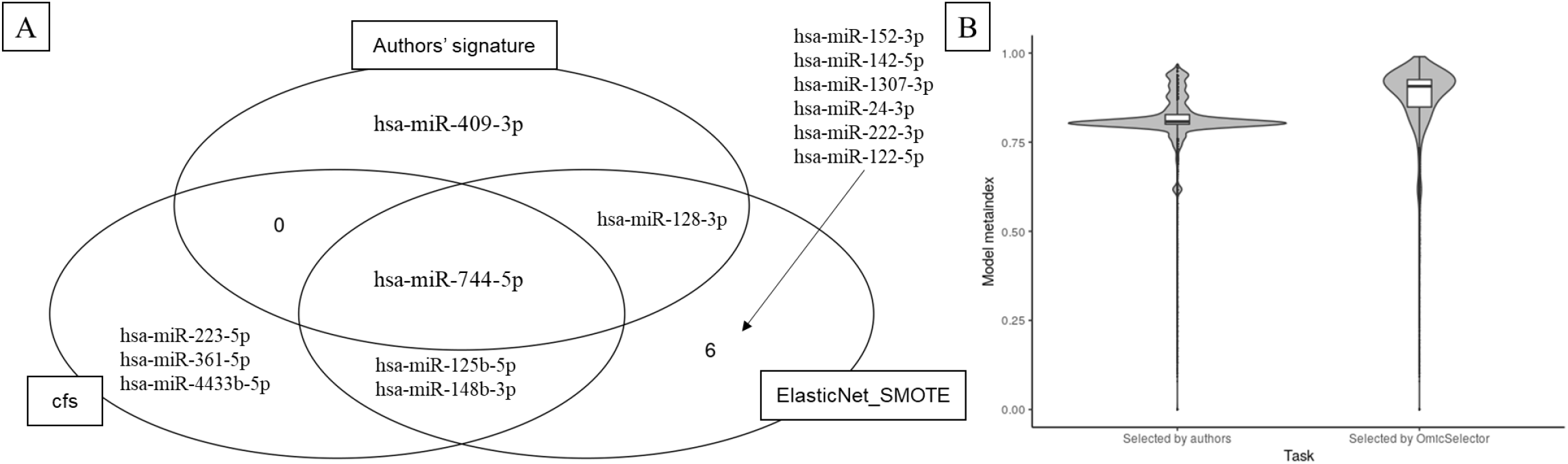
Best signatures overlap and general performance of deep learning OmicSelector full scan. Panel A presents the overlap between the top 2 OmicSelector signatures and the authors’ set [25] of miRNAs. Panel B shows the violin plot comparing deep learning networks’ metaindex values of models inducted on original signature and the best set in OmicSelector analysis selected by the correlation-based feature selection (cfs) method.

Full OmicSelector deep learning scans were conducted on unbalanced training set with “cfs” and original signature. Performance metrics and hyperparameters for each network were provided in Supplementary Table 6. In the OmicSelector full deep learning scans, the signature selected by OmicSelector (“cfs”) performed significantly better than the original one (p<0.0001, Figure 4B). Multivariable analysis has shown that the OmicSelector signature was associated with the most remarkable boost in neural networks’ performance, even when compensating for diverse hyperparameters in the model. Additionally, this linear mixed-effects model showed increased performance was associated with more neurons in all three layers. This increase was most prominent for the first layer and declined in the second and third. Preprocessing of data with a z-score before modeling provided the second most important benefit in performance.

SELU activation function turned out to outperform RELU and sigmoid functions in all three layers. The only exception was the difference between RELU and sigmoid function in the third layer, where no statistically significant effect was noted. The addition of L1 regularization in the first layer improved the performance, but, interestingly, the dropout of neurons lowered it. Lastly, the ADAM optimizer was significantly better than RMSprop and SGD. Coefficients and their 95%CI were presented in Figure 5A and Table 3.

**Figure 5.**
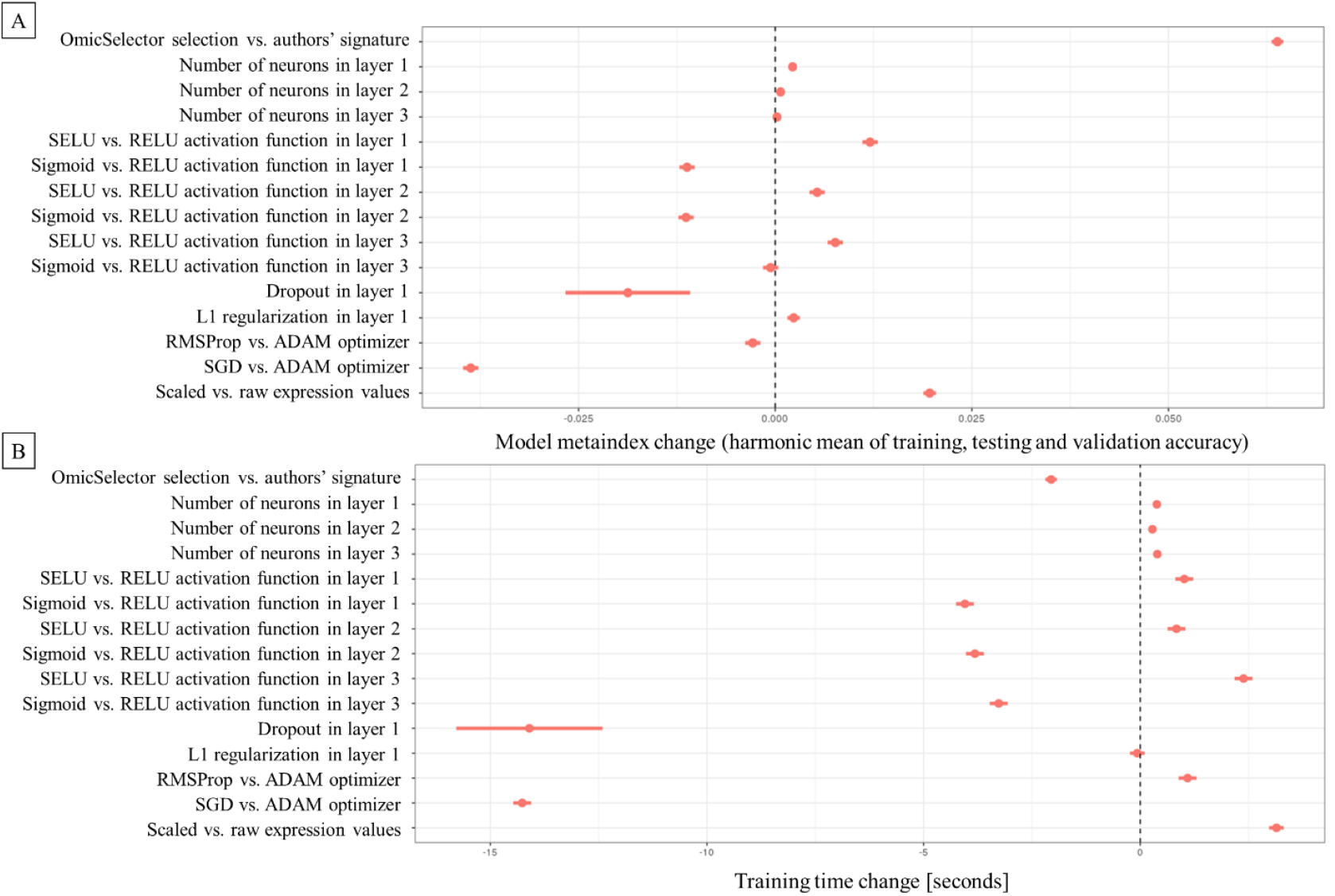
Estimates and their 95%CI for predictors in the linear multivariable mixed-effects model. The figure shows estimates describing the effect of hyperparameters on predictive performance (measured as model metaindex). Panel B presents the estimates relating to hyperparameters’ impact on deep neural networks’ training time.

**Table 3.**
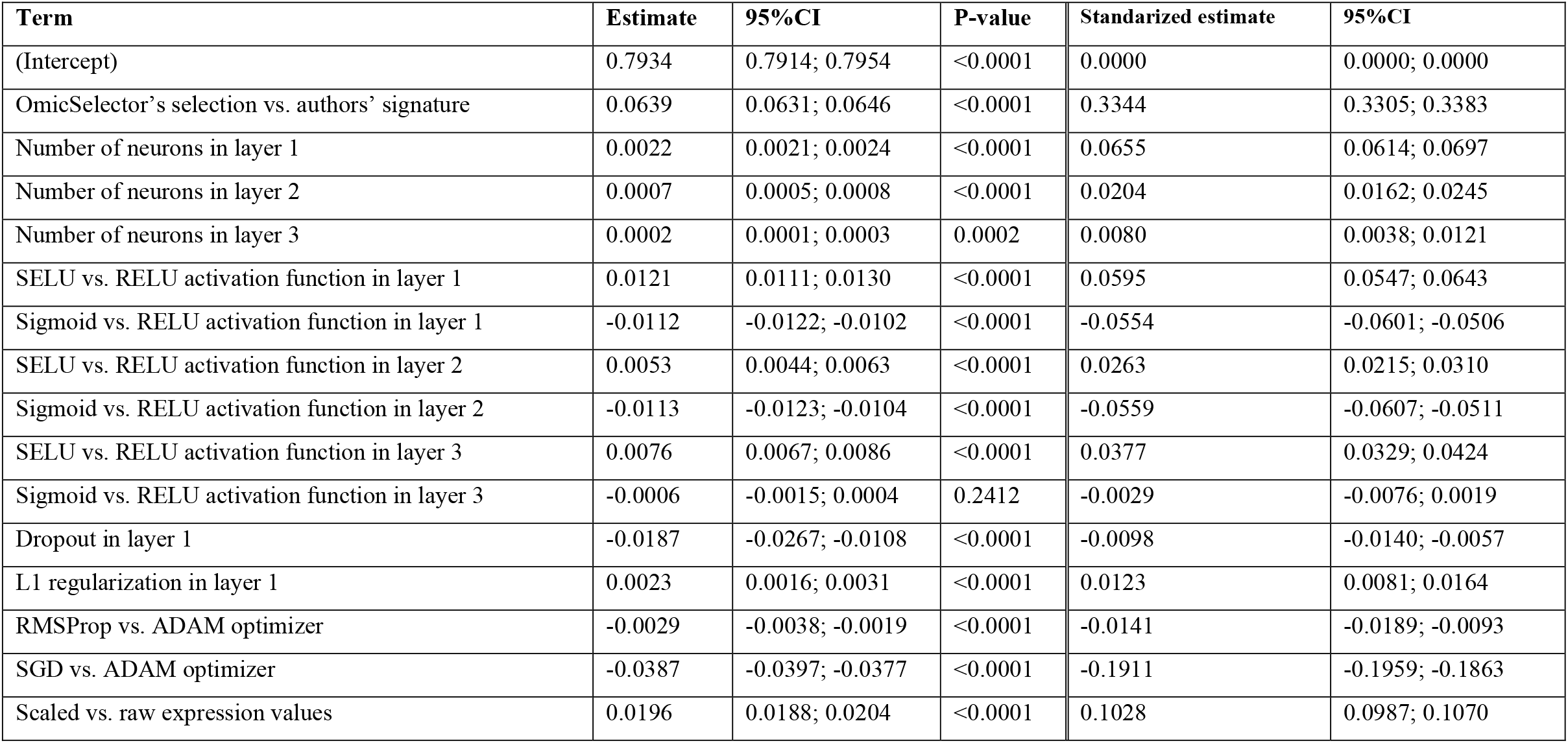
Coefficients of linear mixed-effects model assess hyperparameters’ impact on deep neural networks’ predictive performance (based on model metaindex value as dependent variable). The table presents both crude and standardized estimates.

| Term | Estimate | 95%CI | P-value | Standardized estimate | 95%CI |
| --- | --- | --- | --- | --- | --- |
| (Intercept) | 0.7934 | 0.7914; 0.7954 | <0.0001 | 0.0000 | 0.0000; 0.0000 |
| OmicSelector's selection vs. authors' signature | 0.0639 | 0.0631; 0.0646 | <0.0001 | 0.3344 | 0.3305; 0.3383 |
| Number of neurons in layer 1 | 0.0022 | 0.0021; 0.0024 | <0.0001 | 0.0655 | 0.0614; 0.0697 |
| Number of neurons in layer 2 | 0.0007 | 0.0005; 0.0008 | <0.0001 | 0.0204 | 0.0162; 0.0245 |
| Number of neurons in layer 3 | 0.0002 | 0.0001; 0.0003 | 0.0002 | 0.0080 | 0.0038; 0.0121 |
| SELU vs. RELU activation function in layer 1 | 0.0121 | 0.0111; 0.0130 | <0.0001 | 0.0595 | 0.0547; 0.0643 |
| Sigmoid vs. RELU activation function in layer 1 | -0.0112 | -0.0122; -0.0102 | <0.0001 | -0.0554 | -0.0601; -0.0506 |
| SELU vs. RELU activation function in layer 2 | 0.0053 | 0.0044; 0.0063 | <0.0001 | 0.0263 | 0.0215; 0.0310 |
| Sigmoid vs. RELU activation function in layer 2 | -0.0113 | -0.0123; -0.0104 | <0.0001 | -0.0559 | -0.0607; -0.0511 |
| SELU vs. RELU activation function in layer 3 | 0.0076 | 0.0067; 0.0086 | <0.0001 | 0.0377 | 0.0329; 0.0424 |
| Sigmoid vs. RELU activation function in layer 3 | -0.0006 | -0.0015; 0.0004 | 0.2412 | -0.0029 | -0.0076; 0.0019 |
| Dropout in layer 1 | -0.0187 | -0.0267; -0.0108 | <0.0001 | -0.0098 | -0.0140; -0.0057 |
| L1 regularization in layer 1 | 0.0023 | 0.0016; 0.0031 | <0.0001 | 0.0123 | 0.0081; 0.0164 |
| RMSProp vs. ADAM optimizer | -0.0029 | -0.0038; -0.0019 | <0.0001 | -0.0141 | -0.0189; -0.0093 |
| SGD vs. ADAM optimizer | -0.0387 | -0.0397; -0.0377 | <0.0001 | -0.1911 | -0.1959; -0.1863 |
| Scaled vs. raw expression values | 0.0196 | 0.0188; 0.0204 | <0.0001 | 0.1028 | 0.0987; 0.1070 |

The best neural network (hyperparameter_id of 73814; Supplementary Table 6) presented with the area under the ROC curve of 0.99 (95%CI: 0.96-1.00). This 3-layer model shown an accuracy of 97.06% on the training set, with 100% sensitivity and 92.3% specificity. Its predictions were perfect (100% accuracy) on both testing and validation sets. For reference, the best neural network developed on the authors’ signature (hyperparameter_id of 1005) achieved perfect discrimination on training and validation sets but 90.91% accuracy on the testing set (85.71% sensitivity, 100% specificity).

The average time of one neural network training was estimated as 21.4±19.1 seconds. By massive parallelization of the training process, a “full” deep learning scan using OmicSelector was finished within 29.1 hours on our computing server (Intel(R) Xeon(R) Gold 5120, 256 GB RAM, Nvidia Quadro P6000; training 40 networks in parallel). The relationship between particular hyperparameters and training time is presented in Figure 5B. Screenshot of the final OmicSelector analysis report is attached as Supplementary Appendix.

## 4. Discussion

The developed software – OmicSelector and its corresponding R package, environment, and Docker-based visual application for both biomarker signature selection (feature selection on high-throughput experiment data) and deep learning model development are very powerful and easy-to-use tools for biomarker studies. Our solution’s main advantage is integrating various R- and python-based tools into the reproducible, open-source pipeline for preprocessing, feature selection, and model development. By incorporating a graphical user interface (GUI), our solution is easy to use for users with no or minimal coding skills.

Several solutions for feature selection were previously developed and published; the effort has been recently described in the review by Remeseiro et al. [26]. To the best of our knowledge, our package covers all of the typically used feature selection methods. We publish the R package and the application with GUI, allowing less experienced users to utilize the full potential of complex procedures (like cutting-edge deep learning or genetic algorithms). GUI functionality delivers the whole pipeline and tools to all users and requires no coding or OmicSelector’s function. However, users familiar with R can easily extend both the software and GUI with additional methods. The Docker image provides a pre-configured environment with preinstalled GPU- or CPU-based Keras (with TensorFlow) [27], caret [28], and several helpful R packages. Integrated RStudio, Jupyter-based notebooks, and VS Code offer an ultimate and flexible integrated development environment for further tuning. The software is open-source and highly customizable. Our approach is coherent with the best practices of biomarker development studies due to internal and external validation, thus providing the signature with the most excellent predictive performance and the most outstanding resilience to overfitting. [29]

Several solutions have also been previously published to tackle the automatization of deep learning modeling; [30] however, only a few have considered optimizing the networks for tabular data, as seen in omic experiments. One of the most popular is Auto-Keras [31], which enables Bayesian optimization to guide the network morphism for efficient neural architecture search. However, although complex from the biological standpoint, we described previously that next-generation sequencing expression data could be utilized for correct disease detection by the neural network with just one hidden layer. [3] Therefore, we suggested that the benefit of deep learning in similar problems could be associated with automatic feature extraction and autoencoders achievable by less complex structures. Here, we strictly control hyperparameters not to overcomplicate neural network structure, eventually leading to overfitting. Due to limited complexity, grid search minimizes the impact of chance on the optimization process. Lastly, this tool does not provide GUI.

The presented case study showed that considering OmicSelector’s best signature could significantly boost deep learning performance. The analysis also provided suggestions for the construction of a neural network for serum miRNA-seq data. The final network provided near ideal performance not only on testing but also on the validation set. Due to the small dataset size, we didn’t implement autoencoders (i.e. extensive scan) in the presented case study.

In conclusion, OmicSelector is an easily accessible Docker-based application for biomarker signature selection (feature selection) and deep learning model development based on the results from high-throughput experiments. The software applies 94 feature selection approaches and assesses predictive potential with 11 modeling methods. After the feature set is selected, it performs an extensive grid search to develop the best deep neural network. Less experienced users can use the web-based GUI to perform the analysis, learn the R package or extend the examination with a Jupyter-based notebook, R Studio, or VS Code built-in the application. The Docker-based implementation allows for accessible GPU-based computing to speed up the study of more massive datasets. The tool is open-source and freely available:

- Website, documentation and tutorials: https://biostat.umed.pl/OmicSelector/
- Demo version: https://biostat.umed.pl/OmicSelector/demo/
- Source code: https://github.com/kstawiski/OmicSelector/

## Funding information

The work is supported by National Science Centre in Poland, research grant no. 2018/29/N/NZ5/02422.

### List of Abbreviations

BTCa: biliary tract cancer
CPU: central processing unit
GEO: Gene Expression Omnibus
GPU: graphics processing unit
GUI: graphical user interface
miRNA: microRNA
miRNA-seq: small RNA sequencing
PCa: pancreatic cancer
RELU: Rectified Linear Units
RNA: Ribonucleic acid
RNA-seq: RNA sequencing
ROC: receiver operating curve
ROSE: Random Over-Sampling Examples
SELU: Scaled Exponential Linear Units
SMOTE: Synthetic Minority Oversampling Technique

## Declarations

The study was approved by Bioethical committee at the Medical University of Lodz. Due to the study design, no additional consent was necessary.

## Availability of data and material

All data generated or analysed during this study are included in this published article.

## Authors’ contributions

KS – study design, software development and debugging, case study implementation, manuscript preparation; MK – software development; DM – software testing and improvement suggestions; DM – software testing and improvement suggestions; PH – software testing and improvement suggestions; AD – software testing and improvement suggestions; JS – software testing and improvement suggestions; DC - software testing and improvement suggestions, supervision, critical reading of manuscript; WF - software testing and improvement suggestions, supervision, critical reading of manuscript

## Acknowledgements

Not Applicable

